# Dynamic diffusion analysis of the yeast plasma membrane using Airyscan based microscopic techniques

**DOI:** 10.64898/2026.07.20.739559

**Authors:** Martha S. Cruz-Xelhuantzi, Alex Roof, Blythe Wright, Katherine Paine, Amy Milburn, Grant Calder, Nia Bryant, Peter O’Toole, Ines Hahn, Chris MacDonald

## Abstract

The yeast plasma membrane (PM) is highly compartmentalised into distinct nanoscale domains. The mechanisms by which this organisation regulates surface proteins are not fully understood, and it remains unclear how different biophysical modalities capture diffusion kinetics across varying spatial scales. Using confocal microscopy and an Airyscan2 detector, we benchmarked two prominent techniques: Fluorescence Correlation Spectroscopy (FCS) via the Zeiss Dynamics Profiler and Fluorescence Recovery After Photobleaching (FRAP). We quantified the lateral diffusion of three functionally diverse GFP-tagged model proteins: the exocytic t-SNARE Sso2, the lipid-binding protein Pmp3, and the eisosome-associated protein Ycp4. While diffusion coefficients aligned tightly between both modalities for Pmp3 and Ycp4, Sso2 exhibited a stark 14-fold discrepancy, displaying drastically faster local mobility by FCS compared to macroscopic recovery by FRAP. High-resolution 3D Structured Illumination Microscopy (3D-SIM) shows that Sso2 is partitioned into regional subdomains, that occupy less PM area than the network-like localisation of Pmp3. Our findings suggest that FCS captures rapid, localised diffusion within these microenvironments, whereas FRAP measures highly restricted transit across domain boundaries. Ultimately, this work demonstrates that membrane diffusion coefficients cannot be interpreted in isolation and capturing true lateral mobility requires pairing kinetic measurements with super-resolution spatial mapping to decode complex membrane compartmentalisation.

## INTRODUCTION

As the primary interface between a eukaryotic cell and its environment, the plasma membrane (PM) dynamically aligns a vast array of surface proteins that mediate critical cellular processes. The PM of *Saccharomyces cerevisiae* is no longer viewed as a homogenous fluid mosaic, but rather as a highly ordered patchwork of distinct protein and lipid micro-domains^1,2^. Many years ago, examples of specific subdomains, like the Membrane Compartment containing Pma1 (MCP) and the Membrane Compartment containing Can1 (MCC) were defined^3,4^. The MCP is characterised by the surface membrane H^+^-ATPases Pma1, which functions as a hexamer to maintain cytosolic pH and plasma membrane potential^5,6^. The MCC, also known as eisosomes, correspond to invaginations of the PM that house amnio acid transporters^7,8^, and are associated with physiological stress responses^9–11^. Since then, intricate levels of regulation across PM domains have been demonstrated, with specific proteins being functionally regulated in a spatiotemporal manner between subdomains^12^. As such, the yeast PM represents a sophisticated level of spatial organisation that governs vital cellular processes and serves as a model to study membrane dynamics of eukaryotic proteins. Understanding how proteins navigate this mosaic architecture is essential for deciphering their kinetic regulation, but the precise physical mechanisms that define movement patterns remain incompletely understood.

To quantify lateral dynamics of PM proteins, multiple approaches can be used. Classic examples include Fluorescence Correlation Spectroscopy (FCS) and Fluorescence Recovery After Photobleaching (FRAP), which provide microscopic and macroscopic views of molecular mobility, respectively^13^. While these techniques are frequently used to derive diffusion coefficients, they sample the membrane at fundamentally different spatial and temporal scales: FCS measures high-frequency fluctuations of individual molecules within a diffraction-limited observation volume, while FRAP typically characterises bulk ensemble movement over several micrometers^14,15^. Previous benchmarking studies proved that structural complexities in membranes cause standard confocal FCS and FRAP diffusion estimates to disagree but may have lacked the spatial resolution to define precise patterning^16^.

Advancements in detector technology, such as the Airyscan2 system, offer an opportunity to bridge the gap between diffraction-limited confocal imaging and super-resolution dynamics, for both FCS and FRAP^17– 19^.cA key methodological benefit of Airyscan2 FCS is the capacity to analyse cells exhibiting protein expression levels identical to those used in FRAP studies^20^; standard confocal FCS typically demands sparse fluorophore concentrations to prevent signal saturation, restricting its application to low level proteins^21,22^. By bypassing this limitation, the Airyscan2 detector allows both kinetic approaches to be deployed on a single imaging platform^19^, minimising instrumental variance for improved comparative analysis. High resolution imaging approaches have been used to understand the organisation and dynamics of PM proteins in mammalian cells^23^. However, budding yeast cells are much smaller, and their PM is highly curved, and intensely compartmentalised^2,24,25^. A landmark study using FRAP established that lateral diffusion in the yeast PM is anomalously slow compared to metazoan cells, providing a kinetic mechanism to maintain cell polarity via endocytic recycling^26^. However, mapping these dynamics in yeast using classic diffraction limited FRAP presents potential issues, with the bleach spot representing a large percentage of the total PM signal, an issue exacerbated by equatorial (centre focussed) confocal imaging forcing lateral dynamics into a 1D profile prone to artifacts. In the complex, highly compartmentalised environment of the yeast PM, the assumption that FCS and FRAP provide interchangeable mobility values remains largely untested through rigorous benchmarking. To address this gap, we set out to conduct a direct comparison of FCS and FRAP dynamics within the yeast PM.

By utilising the Airyscan2-based Dynamics Profiler, it is now possible to perform spot-variation FCS with enhanced signal-to-noise ratios and superior spatial precision^17^. In this study, we analysed three yeast membrane proteins to evaluate the consistency of Airyscan2-based FCS and FRAP measurements. Our findings revealed a kinetic paradox for one protein, that exhibits rapid intra-domain mobility via FCS but significantly slower macroscopic recovery by FRAP. We provide a model of micro-domain confinement to rationalise this discrepancy and highlight a single-modality approach can be insufficient for characterising complex membrane architectures and protein dynamics.

## RESULTS

### Characterisation of model yeast plasma membrane proteins

We selected three plasma membrane (PM) proteins as representatives to measure lateral diffusion at the yeast surface (**Figure 1A**). These include: Sso2, a t-SNARE protein with a C-terminal transmembrane domain that promotes exocytic vesicle fusion^27^; Pmp3, which physically binds specific PM lipids and is responsible for regulating plasma membrane polarisation^28,29^; and Ycp4, a protein localised to eisosomes by palmitoylation and implicated in quiescence maintenance^30,31^. To assess these dynamics, protein expression was driven by the constitutive *NOP1* promoter^32^, which establishes a stable, physiological equilibrium of protein turnover and membrane association to eliminate expression-based bias. To accurately quantify lateral diffusion within a uniform plane, imaging and analysis were strictly confined to the apical or basal cortical regions (**Figure 1B**). Surface localisation of the three model proteins was validated by confocal imaging of GFP-tagged strains either stained with FM4-64 to label vacuoles (**Figure 1C**) or co-expressed with Cit1-mTurquoise2 to label mitochondria (**Figure 1D**).

**Figure 1:**
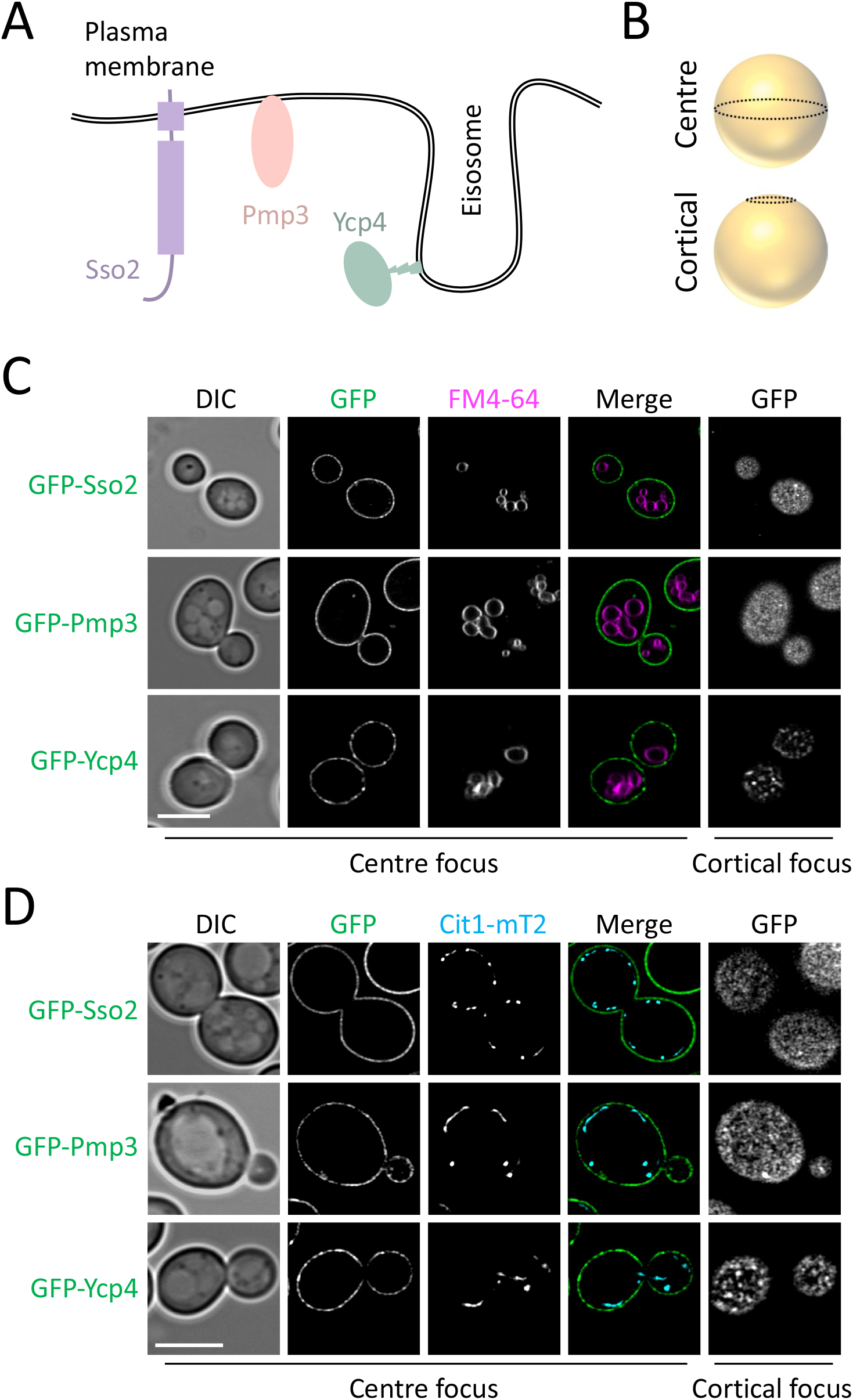
Model yeast proteins for membrane diffusion analysis. **A)** Cartoon representation of three surface localised GFP tagged yeast proteins to serve as models for diffusion analysis. Including the t-SNARE protein, Sso2 (purple), the membrane polarisation factor, Pmp3 (orange), and the eisosome localised factor, Ycp4 (green). **B)** Schematic for centre and cortical focus strategies to image the yeast plasma membrane. **C)** Cells expressing indicated GFP tagged proteins were labelled with a ‘pulse’ of 2µM of FM4-64 in YPD for 30 minutes, washed 3x in SC minimal media followed by a 1 hour ‘chase’ period in label free SC media prior to confocal Airyscan2 microscopy. **D)** Labelled GFP expressing cells were transformed with a plasmid co-expressing Cit1-mTurquoise2 (mT2), grown to mid log phase in selective media followed by Airyscan2 imaging. Scale bar, 5µm

### Diffusion measurements by Dynamics Profiler

Using cortical focussed GFP-Pmp3 as an example, a field of view containing multiple cells was acquired and used to select regions for Dynamics Profiler analysis (**Figure 2A**). The system allows for up to 10 regions to be analysed per run, which can be used if the sample remains stable over the series of acquisitions (3 - 15 seconds per region). To validate the method, we collected baseline tracking data and plotted an autocorrelation function plotted against lag time (**Figure 2B**). The autocorrelation curves were fitted using a 2D diffusion model, rather than a 3D model, to isolate the lateral mobility of the protein within the PM. The characteristic decay profile establishes the baseline mobility of the target protein. For GFP-Pmp3, we confirmed the absence of directional flow and verified isotropic diffusion within the microenvironment. The photostability of GFP-Pmp3 was confirmed across acquisition times (typically 6-10 seconds), with no significant perturbations (e.g. debris or movement through the laser beam) during measurements (**Figure 2C**). These quality control measures were performed for all downstream measurements for the various GFP tagged strains. Employing this technique, we observed distinctive membrane dynamics among the three model proteins. GFP-Sso2 displayed a relatively fast mobility of 0.027 ± 0.011 µm^2^/s (**Figure 2D**). GFP-Pmp3, although significantly slower than GFP-Sso2, also showed relatively fast movement, 0.017 ± 0.003 µm^2^/s. However, eisosomally localised GFP-Ycp4 showed minimal diffusion, averaging 0.003 ± 0.001 µm^2^/s. To ensure these measurements represented the physiologically relevant diffusion across the plasma membrane, measurements were repeated as before for GFP-Sso2 in standard media or following exchange with an azide kill buffer to inhibit ATP production, to stall metabolism and trafficking whilst maintaining spatial architecture^33^. As expected, this had a profound effect on the dynamics of the GFP-Sso2, with a significant decrease from 0.025 ± 0.008 µm^2^/s down to 0.007 ± 0.004 µm^2^/s (**Figure 2E**). This was still slightly higher than the relatively immobile measurements of GFP-Ycp4, 0.004 ± 0.001 µm^2^/s.

**Figure 2:**
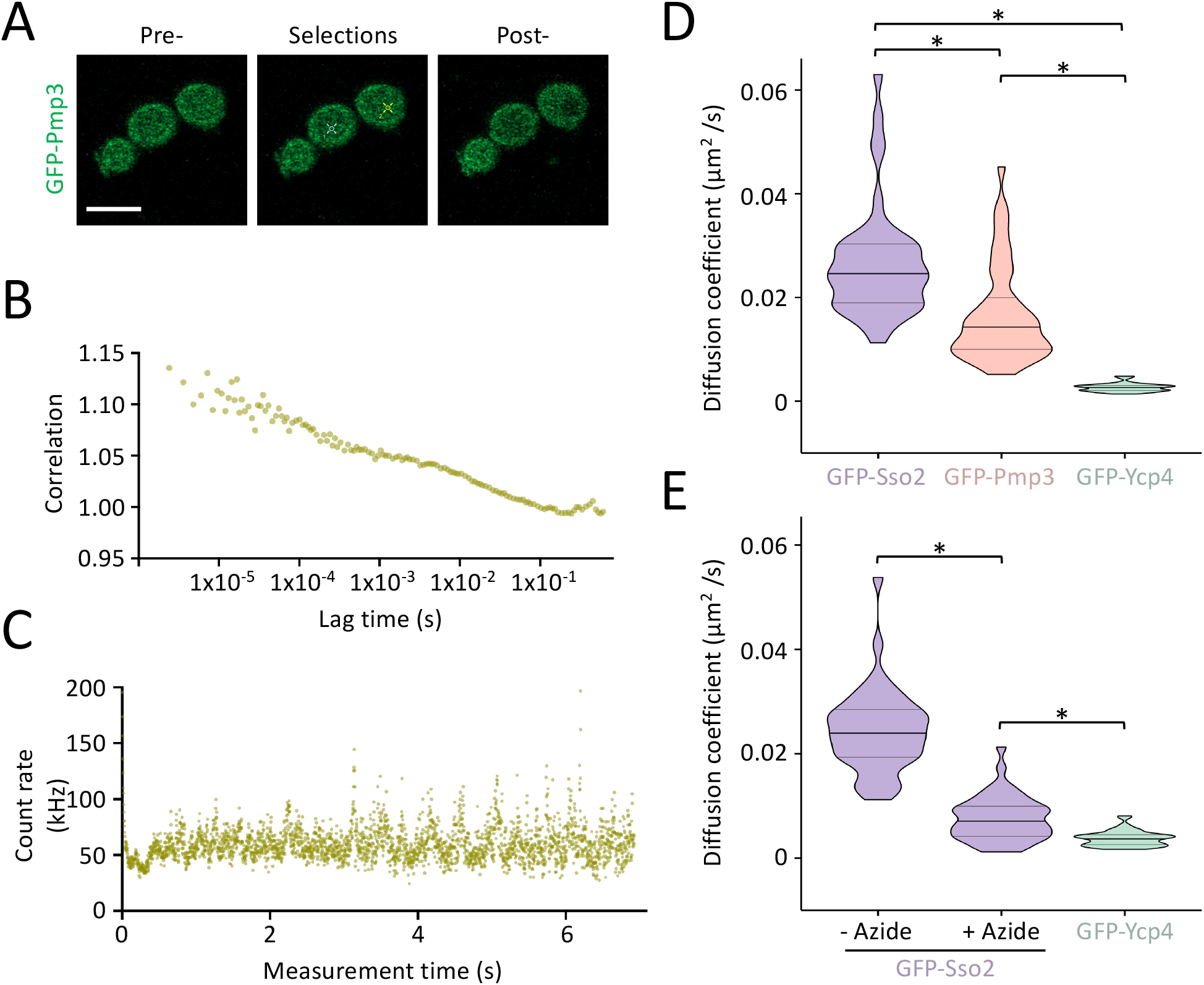
Dynamics Profiler FCS measurements show fast and slow diffusion rates. **(A)**Representative field of view of cells expressing GFP-Pmp3, with the selection of regions (2 shown) for analysis indicated and the pre- and post-fluorescence measure following a 15s Dynamics Profiler acquisition. **(B)**Representative temporal autocorrelation curve of raw collected data points, demonstrating 2D lateral mobility within a single selection of GFP-Pmp3 expressing cells with cortical focus. **C)** The raw photon count rate of representative GFP-Pmp3 acquisition tracked continuously over total measurement time. **D - E)** Mean diffusion co-efficient estimates for each of the indicated GFP tagged proteins under optimal media conditions **(D)** and for GFP-Sso2 following exchange of media with buffer containing azide (**E**). Scale bar, 5µm. Data represent mean standard deviation from biological replicates (n = 3). Statistical significance was determined using a two-tailed, unpaired Student’s *t*-test, * indicates p < 0.001.

### Diffusion validation by FRAP

The diffusion parameters correlated with reasonable values for each of the three sample proteins, given they are localised to the confirmed yeast PM^25,34^. To validate these measurements, we employed FRAP of the same proteins with a similar experimental approach. For this, cortical focussed cells were imaged, with a small region of each cell chosen for bleaching. 20 frames prior to bleach were acquired, the first 10 to stabilise the system and the second 10 to provide a pre-bleach average for downstream calculations. The percentage recovery after bleaching was then monitored. To begin with, we used a relatively short time intervals of 125 ms but this showed little recovery of GFP-Sso2 after 100 acquisitions (**Figure 3A**), in line with previous observations^26^. Using the same protocol, we next analysed cells expressing GFP-Pmp3 and GFP-Ycp4. Both targets replicated the Dynamics Profiler baseline: GFP-Pmp3 recovered relatively rapidly, whereas GFP-Ycp4 showed no recovery across the short-interval protocol (**Figure 3B - 3C**). Because neither GFP-Sso2 nor GFP-Ycp4 recovered during this fast (<15s) acquisition window, we repeated the experiments using a longer 2-second interval for 100 iterations, which allowed sufficient recovery for GFP-Sso2 (**Figure 3D**). Somewhat surprisingly, GFP-Ycp4 also recovered in this period, albeit with a different localisation pattern (**Figure 3E**). While the relative diffusion rates of GFP-Pmp3 and GFP-Ycp4 measured by FRAP were consistent with the Dynamics Profiler results, those for GFP-Sso2 revealed an unexpectedly slow diffusion rate (**Figure 3F**). Consequently, the degree of convergence between the two modalities differed substantially depending on the specific model protein analysed (**Figure 3G**). GFP-Pmp3 and GFP-Ycp4 exhibited tight alignment across respective FCS and FRAP systems: yielding respective values of 0.024 versus 0.017 µm^2^/s, while the immobile GFP-Ycp4 differed by a negligible margin of 0.005 versus 0.003 µm^2^/s. In stark contrast, the measured mass transport of GFP-Sso2 by Dynamics Profiler was 0.027 µm^2^/s, representing a nearly 14-fold increase in diffusion velocity compared to the slow 0.002 µm^2^/s baseline calculated by FRAP.

**Figure 3:**
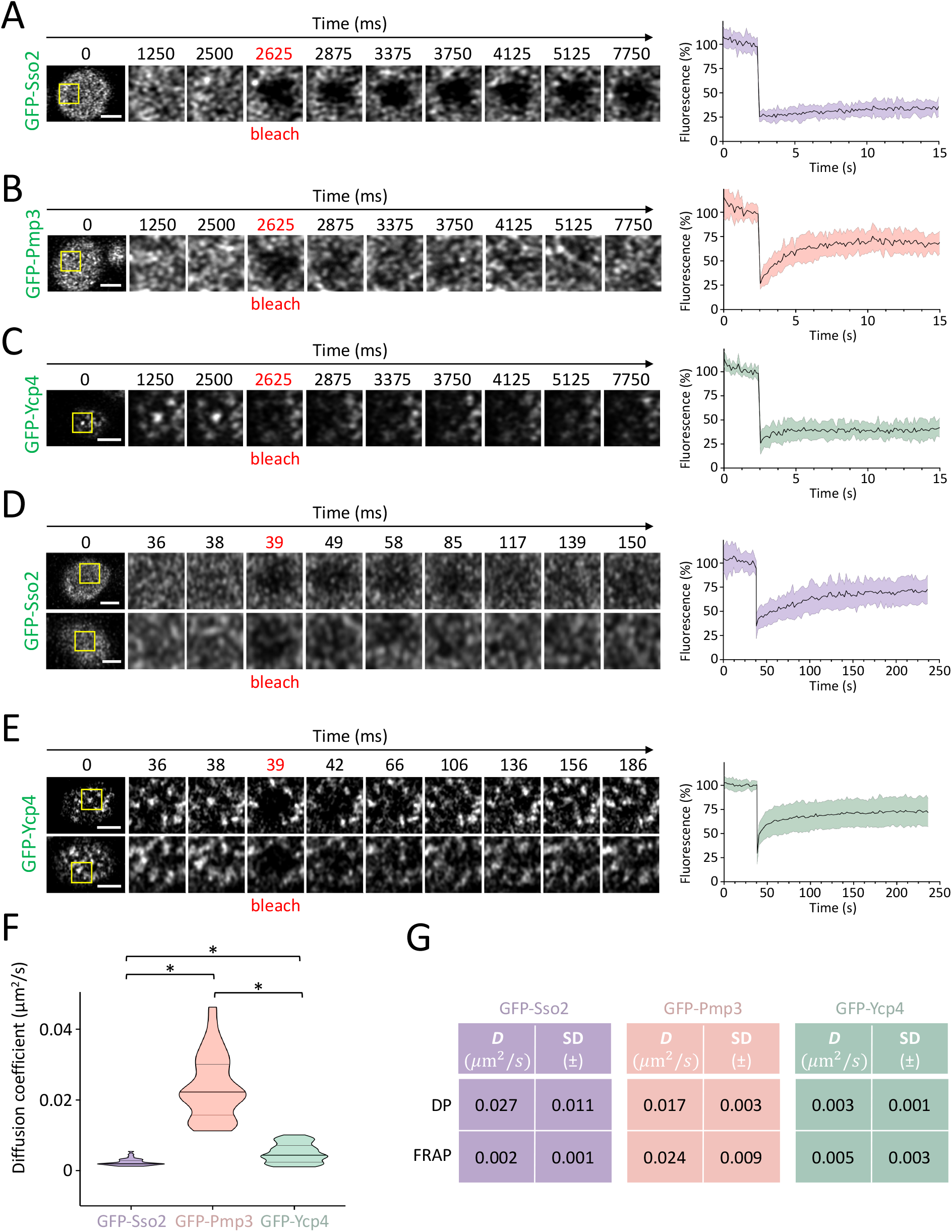
FRAP analysis shows varied diffusion of GFP tagged proteins. **A - E)** Time-lapse series of FRAP experiments were optimised for fast **(A - C)** and slower **(D - E)** recovery of cells expressing: GFP-Sso2 (purple), GFP-Pmp3 (orange), and GFP-Ycp4 (green). Each inset (yellow box) from a representative cell shows examples of surface fluorescence pre- and post-bleach (red). Average recovery profiles (n>30) are plotted in graphs (right) **F)** Diffusion coefficients were calculated for indicated proteins following FRAP analysis. **G)** Tables comparing diffusion coefficient estimates using Dynamics Profiler (DP) and fluorescence recovery after photobleaching (FRAP). Scale bar, 5µm. Data represent mean standard deviation from biological replicates (n = 3). Statistical significance was determined using a two-tailed, unpaired Student’s *t*-test, * indicates p < 0.001.

### Spatiotemporal pattering of the yeast PM

Previous work has shown that individual surface proteins localised to the yeast PM exhibit distinct localisations patterns^2,12^. Consequently, we sought to test whether these spatial profiles could account for the disparate diffusion rates observed between the two measurement systems. We have previously localised the endosomal recycling inhibitor Gpa2 to the surface in a relatively uniform, networked localisation pattern^35^, which is spatially distinct from the eisosme marker Pil1 labelled with mCherry (**Figure 4A**). We confirmed these localisation profiles using multiple high-resolution imaging modalities, standard confocal, Airyscan 2, and Lattice Structured Illumination Microscopy (SIM). Notably, Airyscan2 and Lattice SIM outperformed standard confocal, yielding sufficient spatial resolution to delineate Gpa2-GFP patterns that are spatially segregated from eisosomes (**Figure 4B**). Building on these experiments, we utilised the eisosome marker Pil1-mCherry as a robust spatial landmark to map and characterise the precise localisation patterns of our three model proteins (GFP-Sso2, GFP-Pmp3, and GFP-Ycp4). Using 3D-SIM, we captured high-resolution images across both the equatorial and cortical planes of the cell, generating average intensity projections for each to resolve their spatial distribution relative to eisosomes (**Figure 4C**).

**Figure 4:**
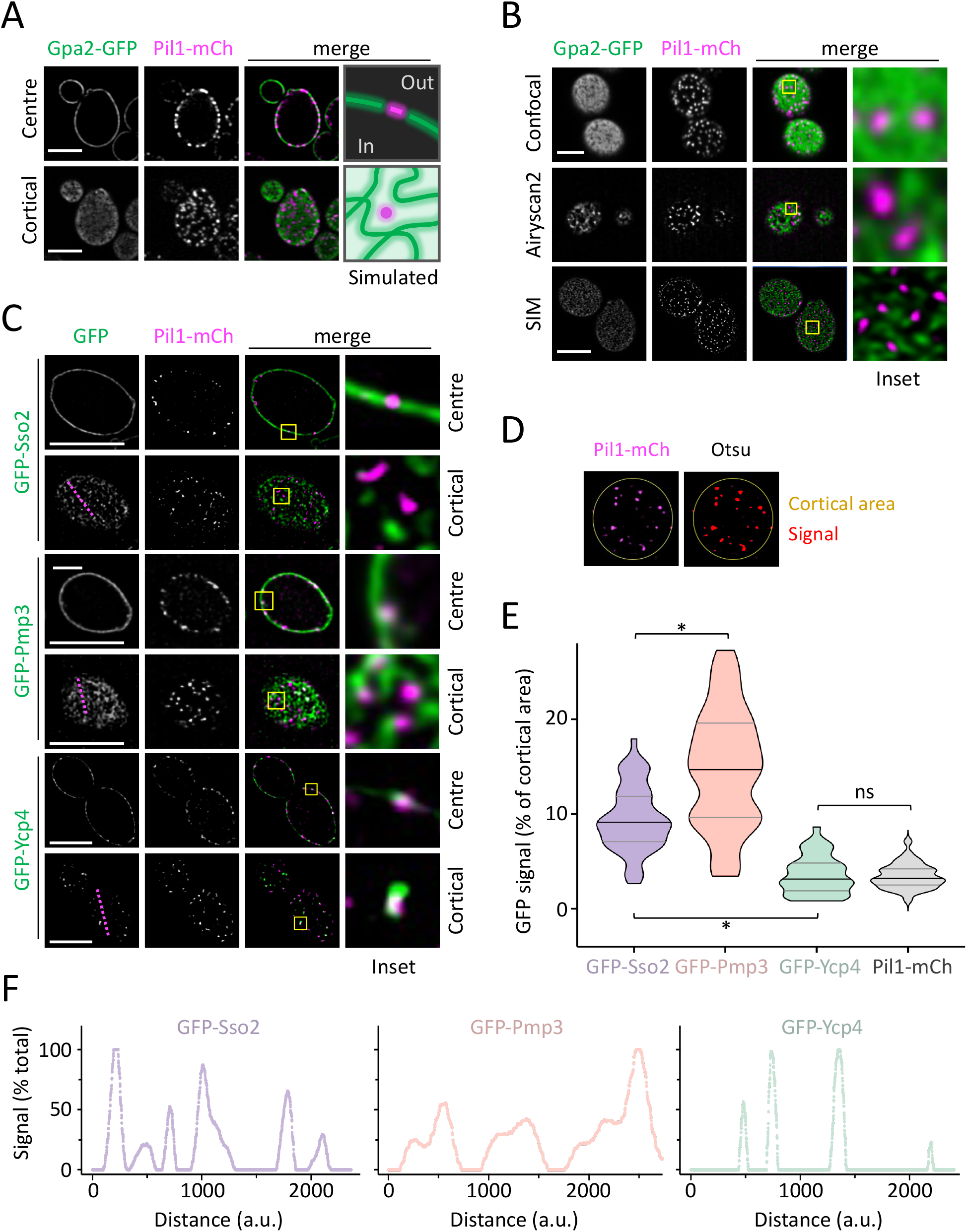
Spatiotemporal patterning of GFP tagged yeast PM proteins. **A - B)** Cells co-expressing Gap2-GFP and Pil1-mCherry were grown to log phase prior to imaging with indicated modalities. **C)** Diploid strains expressing GFP-tagged proteins alongside Pil1-mCherry were imaged using SIM and insets (yellow box) show enlarged regions of interest. **D)** Segmentation strategy to isolate signal from cortical regions. **E)** Analysis of cortical region containing distinct regions or network-like localisations patterns. **F)** Line analysis, magenta in **(C)**, for indicated profiles of GFP-tagged proteins. Scale bar, 5µm. Data represent mean standard deviation from biological replicates (n = 3). Significance determined using a two-tailed, unpaired Student’s *t*-test, * indicates p < 0.001; ns = not significant.

As anticipated, GFP-Ycp4 co-localised with Pil1-mCherry within low-mobility eisosome compartments^36,37^. Conversely, both GFP-Sso2 and GFP-Pmp3 exhibited distinct regional localisation profiles with prominent subdomain architectures across the PM. To quantify these distribution patterns, we segmented cortical regions and calculated the percentage area occupied by GFP (**Figure 4D**). GFP-Sso2 and GFP-Pmp3 differed significantly; the latter presented an interconnected, network-like pattern that accounted for a greater proportion of the total membrane surface (**Figure 4E**). This model was supported by line-profile analysis of representative images, demonstrating that GFP-Sso2 localises to more discrete regions than GFP-Pmp3, while still occupying a larger fraction of the membrane surface than eisosomes marked by GFP-Ycp4 (**Figure 4F**). We hypothesise that while Sso2 diffusion remains highly rapid within individual, isolated microdomains detected by the Dynamics Profiler, long-range transit between these distinct pools is constrained. This suggests that molecules face physical or energetic barriers when navigating intervening subdomains that contain negligible steady-state levels of GFP-Sso2, thereby accounting for the slow macro-diffusion observed by FRAP.

## DISCUSSION

In this study, we have established and optimised a robust methodology for benchmarking lateral protein diffusion in the compartmentalised plasma membrane (PM) of *Saccharomyces cerevisiae*. By pairing a high-resolution, Airyscan2 approach to FCS-based Dynamics Profiler and Fluorescence Recovery After Photobleaching (FRAP), we have demonstrated that membrane mobility dynamics cannot be fully understood through a single kinetic assessment alone. While a highly restricted structural marker and a uniformly distributed proteolipid showed excellent kinetic alignment between FCS and FRAP, an exocytic t-SNARE revealed a striking paradox being fast by FCS and decidedly slow by FRAP. By combining these kinetic approaches with super-resolution 3D-SIM spatial mapping, we indicate that this kinetic discrepancy might be driven by physical domain compartmentalisation. Ultimately, this work provides a practical workflow and a cautionary framework, demonstrating that membrane diffusion coefficients cannot be interpreted in isolation without high-resolution spatial mapping.

To ensure this benchmarking framework was technically accurate, we first optimised the focus strategy and validated FCS measurements for the 3 distinct model GFP tagged proteins: Sso2, Pmp3 and Ycp4 (**Figure 2**). To confirm our protocol accurately captures biologically active membrane dynamics rather than passive physical artifacts, we used metabolic inhibition of cells expressing fast moving GFP-Sso2. We noted that although treating GFP-Sso2-expressing cells with sodium azide massively reduced diffusion, the residual mobility measurements remained higher than those of the GFP-Ycp4 negative control. We assume the diffusion observed in the presence of azide might be explained by incomplete inhibition during acute treatment, or by passive thermal fluctuations^38,39^. Furthermore, even when Sec18 activity is inhibited, post-fusion SNARE interactions within the PM, or other clustering events, might account for some remaining mobility^40,41^. Despite this, the greatly reduced diffusion estimates of GFP-Sso2 following azide treatment suggest that our approach is accurately assessing the mobility of surface molecules.

By employing the eisosome-associated protein GFP-Ycp4 as a static control, both the Dynamics Profiler and our fast-interval FRAP protocol successfully confirmed its highly restricted baseline mobility. This aligned with expectations, as Ycp4 is a stress response factor structurally anchored to the cytoplasmic leaflet of the eisosome via palmytoilation^29,31,42^. Intriguingly, extending the FRAP acquisition window captured a slow fluorescence recovery profile for Ycp4, yet the recovering signal completely lacked the characteristic punctate architecture of eisosomes (**Figure 3E**). Instead, the recovery appeared diffuse across the cortical plane, suggesting that it may not represent genuine lateral diffusion of membrane-bound eisosomal Ycp4. Rather, we reason that this protocol captures a transient cytosolic pool re-decorating the membrane, or perhaps a gradual, de novo accumulation at the eisosome. While Ycp4 represents a structurally locked eisosomal marker, other membrane proteins present a more complex spatial dynamic. We note that, unlike GFP-Sso2, which is entirely distinct from eisosomes, a pool of GFP-Pmp3 localises to both eisosomes and distinct surface domains, which were well defined by 3D-SIM (**Figure 4C**). Strains lacking *PMP3* have been shown to have defects in hyperpolarisation of the PM and reduced levels of Pil1 at eisosomes^29^. It may be that these distinct pools of Pmp3 exhibit varying degrees of diffusion.

While structurally uniform or highly restricted proteins show excellent alignment between FCS and FRAP measurements, the exocytic t-SNARE Sso2 reveals a striking contrast. This kinetic discrepancy by FRAP can perhaps be rationalised by the distinct biological locations shown by 3D-SIM, as well as the functions of these proteins; Sso2 is likely functionally constrained to specific regions of exocytosis or downstream recycling^43,44^, unlike Pmp3, which has broader access to PM regions to mediate functionals related to stress sensing ^28,29^. Indeed, our 3D-SIM data demonstrate that Sso2 exhibits discrete regionality, being partitioned more tightly than the broad, network-like mesh of Pmp3. That said, GFP-Sso2 remained less restricted than the punctate distribution of eisosomes, and there was no overlap between even the punctate localisations of GFP-Sso2 and eisosomes, as shown by co-localisation using Pil1-mCherry (**Figure 4C**). We quantified this spatial patterning using two independent segmentation and line-profile approaches (**Figure 4D – 4F**). These observations are in line with fixed-cell super-resolution microscopy of the organisation of yeast surface proteins, which has shown that proteins like Pma1 and Mup1 are tightly organised^45^.

Curiously, even when examining these established model proteins, caution is required when interpreting comparative data. A case in point is Sso2, which shares 72% sequence identity with its paralogue, Sso1. While their individual deletion yields no obvious phenotypic impact, their simultaneous deletion is lethal^27^. Previous studies have demonstrated that Sso1 and Sso2 do not localise with the same degree of symmetry between mother and daughter cells, a disparity in polarisation revealed through both proteomic and microscopic techniques^46^. This suggests that distinct regulatory mechanisms control the surface dynamics of these highly homologous proteins, a divergence that might be anticipated given specialised function of Sso1 during sporulation^47^.

While this study focused on three model proteins, our workflow serves as a highly adaptable methodological framework that can be readily scaled to encompass any membrane cargo. Far from being a limitation, these three distinct proteins represent diverse archetypes of membrane dynamics, demonstrating that our dual-method approach can resolve even highly complex spatiotemporal profiles. Crucially, the utility of pairing high-resolution fluctuation spectroscopy with super-resolution spatial mapping extends far beyond budding yeast. This integrated pipeline is directly applicable to other challenging, highly compartmentalised biological systems, ranging from: the densely organised cytoskeleton^48^ and receptor trafficking at neuronal synapses^49^ of certain mammalian cells, to the rigid, microdomain-rich envelopes of plant cells^50^ and bacteria^51^. Ultimately, as commercial detector technologies such as Airyscan2 advance accessible biophysics^52^, moving beyond single-modality kinetic measurements will be essential to truly decode how the spatial architecture of living membranes governs molecular function.

## METHODS

### Reagents

Yeast strains and plasmids used in this study are detailed in Supplemental Table T1.

### Cell culture

*S. cerevisiae* yeast strains were grown in either yeast extract peptone dextrose (YPD) media (1% yeast extract, 2% peptone, 2% dextrose) or synthetic complete (SC) minimal media (2% glucose, 0.675% yeast nitrogen base without amino acids, supplemented with base/amino acid drop-out for selections) (Formedium, Norfolk, UK). Cells were typically harvested from overnight serial dilutions, grown to mid-log phase at 30°C with shaking, before preparing for imaging the following morning. Expression of plasmids containing the *CUP1* promoter was induced by addition of 100 µM CuSO_4_ to the media and grown overnight to mid-log phase before harvesting. The azide kill buffer previously described^53^ contains 10 mM sodium azide and 10 mM sodium fluoride, in addition to 50 mM Tris.HCl pH 8.0.

### Yeast mating

To generate diploid strains for co-localisation analysis, *MAT*a strains expressing the respective GFP-tagged model proteins were mated with a *MAT*α strain expressing Pil1-mCherry^11^. Diploids were selected on SC media lacking uracil and histidine, providing dual selection for *URA3*-GFP tagged Sso2, Pmp3, and Ycp4 in addition to Pil1-mCherry-*his5*^*+*^.

### Confocal microscopy

Yeast cells expressing GFP-tagged proteins were grown to mid-loge phase (unless otherwise stated) and concentrated in SC media before imaging. Cells were loaded to a glass slide, and microscopy was performed with a Zeiss 980 (equipped with Airyscan2 detector) laser-scanning confocal instrument using a 63x/1.4 objective lens. Green, red, and blue fluorescence was excited using excitation wavelengths of 488 nm, 561 nm, and 405 nm, respectively, with emissions measured at wavelengths of 495/500 nm, 570/620 nm and 460/550 nm, respectively. Images were processed using Zen Blue (Zeiss) standard Airyscan2 algorithm and modified for publication using ImageJ software (NIH). Yeast vacuoles were labelled with YPD media containing 2µM FM4-64 followed by 3 washes and a 1-hour chase period in SC media prior to imaging.

### Structured illumination microscopy

Super-resolution imaging of the yeast PM was performed using a ZEISS Elyra 7 equipped with Lattice SIM imaging mode (Zeiss, Germany) and an Plan-Apochromat 63x / 1.4 oil objective. GFP was excited at 488nm and its emission was collected 500-550nm and mCherry was excited at 561nm and its emission collected 570-630nm. Phase steps and rotation angles were automatically set by the software to optimize spatial resolution in 2D/3D. Z-stacks of whole yeast cells were collected with 100nm spacing. All raw structured illumination datasets were processed using the SIM^2^ reconstruction algorithm integrated within the Zen Blue software, as previously described^35^. Following reconstruction, channel alignment was performed using a calibration map generated from multispectral fluorescent bead standards to correct for chromatic aberration. Final image analysis, localized crop generation, and brightness/contrast adjustments were carried out using Zen Blue and/or Fiji/ImageJ (NIH, Bethesda).

### Dynamics Profiler

Dynamic Profiler measurements were taken with a Zeiss 980 system (Carl Zeiss Microscopy, Germany) equipped with a Apochromat 40x / 1.2 water objective. Excitation was achieved using a 488nm laser focused directly onto the plane of the yeast cell cortex. The Dynamic Profiler wizard allow for Pre and Post Airyscan2 image collection for ROI targeting and sample assessment. Autocorrelation curves were generated from the intensity fluctuations and fitted using a 2D-diffusion model within the Zen Blue Dynamics Profiler software to calculate lateral diffusion coefficients. To minimise spatial artifacts, only flat, well-defined regions of the peripheral PM were selected for profiling.

### FRAP acquisition

Fluorescence recovery after photobleaching (FRAP) experiments were performed with a Zeiss 980 (equipped with Airyscan2 detector) using a 63x/1.4 objective lens. To establish a baseline, a pre-bleach sequence was recorded consisting of a 10-frame stabilisation period followed by 10 frames to calculate the initial pre-bleach intensity. A circular region of interest (ROI) of 0.7 µm in diameter was then photobleached using a high-intensity laser pulse. Post-bleach recovery was monitored by capturing 100 consecutive frames to track the fluorescence signal return. Images were taken every 125 ms in short protocols and every 2 s in long protocols. Recoveries were recorded 100 post-bleach frames, in both short and long protocols. FRAP evaluations were performed with Zen Blue.

### FRAP analysis

Raw intensity data within the bleached ROI were extracted and analysed. The mobile fraction was calculated as the ratio of recovered fluorescence (plateau minus post-bleach minimum) to total bleachable fluorescence (pre-bleach average minus post-bleach minimum). The half-time of recovery *t*_1/2_ was graphically identified for each dataset, as the time taken to reach the midpoint between the post-bleach minimum and the recovery plateau. This value was then used alongside the squared bleach radius (*w*^2^) to calculate the lateral diffusion coefficient via the Soumpasis equation.

## ACKNOWLEDGMENTS

We would like to thank staff at the York Bioscience Technology Facility for technical assistance. This research was supported by a Sir Henry Dale Research Fellowship from the Wellcome Trust and the Royal Society 204636/Z/16/Z (CM).

